# Cellular heterogeneity in hypertrophic burn scars in response to carbon dioxide laser therapy

**DOI:** 10.1101/2024.07.05.600800

**Authors:** Yung-Yi Chen, Christopher Mahony, Jason Turner, Charlotte M Smith, Abdulrazak Abdulsalam, Ezekwe Amirize, Amberley Prince, Adrian Heagerty, Claudia Roberts, Adam Croft, Yvonne Wilson, Naiem Moiemen, Janet M Lord

## Abstract

Fractional carbon dioxide (AFCO_2_) laser therapy is used for treating pathological scarring, but the clinical outcomes are variable and the mechanisms of scar reduction poorly understood. We investigated the mechanisms underpinning efficacy of AFCO_2_ laser therapy, performing single-cell RNA sequencing in skin biopsies from patients with hypertrophic scars after AFCO_2_ laser therapy. Patients with younger scars (Good Responder, GR, <6 years from healing) had better scar reduction than patients with older scars (Poor Responder, PR, >6 years from healing) by various measures of scarring. scRNAseq analysis revealed that genes enriched in GR were associated with extracellular matrix and structure organisation (*COL14A1*, *POSTN*, *SPARC*); whereas genes enriched in PR were related to enhanced immune responses (*IL-12*, *MSTN, HLA-DQA*). The groups had distinct intercellular communication networks and differentiation trajectories after AFCO_2_, with regenerative Mesenchymal fibroblasts associated with a good response and inflammatory Secretory Papillary and Inflammatory Fibroblasts with a poor response.

## Introduction

Burn injury is one of the most common traumas worldwide with an estimated 11 million people requiring medical attention each year^1^. In the UK alone, approximately 250,000 people with burn injuries present to primary care each year and a further 175,000 patients present to emergency departments. The impact of burn injuries is clinically significant because they often give rise to exuberant scarring that results in permanent physical function loss and psychological issues due to the stigma of disfigurement. Scarring post burn occurs in 32-72% of patients^2^ and more than 100,000 patients develop problematic hypertrophic scarring (HTS) every year in the UK.

In clinical practice, several modalities are commonly used to treat post-burn HTS, such as topical emollients, compression garments, steroid injection or the use of anti-neoplastic drugs such as fluorouracil and bleomycin^3–6^. Laser therapy developed in the 1980s and has been used for treating HTS with variable positive outcomes and minimum side effects^7, 8^. There are three main types of laser therapy being used for treating established scars^9–12^. The 585nm pulsed-dye lasers (PDL) are used to selectively target blood vessels in the scar tissue and reduce scar erythema^8, 13^. Q-switched neodymium-yttrium aluminum-garnet (Q-switched Nd:YAG) lasers emit light with a wavelength of 1064nm and have deeper tissue penetration, which can induce wound healing through a thermal effect on the dermis with minimum effect on the epidermis^14^. AFCO_2_ laser therapy has been shown to induce haemostasis and stimulate fibroblasts to produce new collagens^9, 14–18^. It functions through emitting a wavelength of 10,600nm laser beam, whose energy is quickly absorbed by water which then evaporates to induce tissue damage and break down any scarring matrix.

Clinical evidence has shown that AFCO_2_ treatment is an effective way to treat HTS. For example, in a prospective study, Miletta et al., showed that AFCO_2_ laser therapy provides significant, sustained improvement of elasticity, thickness, appearance and symptoms of mature HTS in adults^19^. Patel et al., further demonstrated that AFCO_2_ laser therapy can improve hypertrophic burn scars in paediatric patients^20^. However, despite being an effective approach to alleviate HTS with positive results overall, clinical outcomes remain variable and the detailed mechanisms of action behind AFCO_2_ laser therapy remain unclear^21^.

Wound healing is a multifactorial, long-term biological process involving a sophisticated dynamic balance within the skin microenvironment. Any variation in the biological process can potentially lead to alternative outcomes, such as hypertrophic scarring. Recent studies have shown that dermal fibroblasts are highly heterogeneous post-wounding and the diversity of fibroblasts at the wound site contributes to skin repair and regeneration, which ultimately impact on scarring outcomes^22–24^. One fibroblast sub-type, the engrailed 1 (*En1*) expressing cells were found to be associated with scarring in mice^25^. Therefore, in this study we aimed to further investigate the complex biological mechanisms underlying the laser therapy to reduce hypertrophic scarring, specifically the cellular infiltrate at the treated scar site, that may contribute to the variable scar reduction outcomes after AFCO_2_ laser therapy.

## Results

### Participant demographics

14 adults aged between 20 and 70 years of age (9 males, 5 females; age 46.57±4.23 years) with established hypertrophic burn scars and who met the inclusion criteria^26^ were recruited to the study. All patients had scars resulting from deep or full thickness burns with TBSA (Total Body Surface Area) ranging from 10 to 85% (average 50.31±7.25%). All the burns were caused by flame and the site of injury included arm (n=2), chest (n=4), back (n=3), thigh (n=4) and prescapular (n=1) areas. The age of the scars, defined as the time from wound closure to the time of AFCO_2_ treatment, ranged from 2 to 65 years (average 21.71±6.45 years), Table 1.

**Table 1.** Participant demographics.

| Participants | Total (n=14) | Young Scar (n=7) | Old Scar (n=7) |
| --- | --- | --- | --- |
| Scar age (years) | 21.71±6.45 (2-65) | 3.57±0.34 (2-6) | 39.86±5.69 (18-65) |
| Participant Gender (M/F) | 9/5 | 4/3 | 5/2 |
| Participant Age (years) | 46.57±4.23 (20-70) | 40.14±4.02 (22-59) | 53±4.84 (27-70) |
| Body Mass Index (BMI) | 26.08±2.65 (20.93-40.52) | 28.33±1.92 (21.55-40.52) | 23.83±3.26 (20.93-34.97) |
| Fitzpatrick Skin Type | 2-4 | 2-4 | 2-4 |
| % TBSA | 50.31%±7.25% (10%-85%) | 53.28±7.16 (10-85) | 46.83±7.32 (17-77) |
| Co-mobility (Y/N) | 4/11 | 2/7 | 1/7 |
| Biological site |  |  |  |
| Arm | 2 | 1 | 0 |
| Chest | 4 | 2 | 2 |
| Back | 3 | 1 | 1 |
| Thigh | 4 | 2 | 0 |
| Prescapular | 1 | 0 | 1 |

### Laser treatment is more effective on less established scars

To investigate the impact of AFCO_2_ laser therapy on hypertrophic burn scars, all participants were given 3 sessions of AFCO_2_ laser treatment at 3 month intervals. Before each treatment and at the end point at year 1 (Y1), the physical characteristics of the scars were examined using different assessment tools, including quantitative scar assessment devices, such as DermaScan® Ultrasound, DSMIII® colourimeter; cutometer; and qualitative scar assessment questionnaires, such as mVSS and POSAS-O. Overall, AFCO_2_ laser therapy resulted in significant colour shift ( Δ E) between month 3 (M3) and Y1 endpoint (5.93±0.76 vs. 19.64±2.70, p=0.0005) and between M6 and Y1 (8.82±2.31 vs. 19.64±2.70, p<0.005). It also produced a significant colorimetric change from baseline to endpoint at Y1 for lightness (43.29±2.88 vs. 59.49±2.43; p<0.0001) and yellowness (11.04±1.10 vs. 8.39±1.46; p<0.05), Table 2 and Supplementary Figure S1. However, other scar assessments between baseline and Y1 endpoint, including mVSS overall score (5.14±0.31 vs. 4.64±0.29), skin thickness (2.64±0.12 mm vs. 2.67±0.19 mm), skin stiffness R0 (0.60±0.07 vs. 0.66±0.08), skin visco-elasticity R2 (0.85±0.02 vs. 0.86±0.02), erythema index (10.88±0.81 vs. 9.27±0.95) and melanin (pigmentation) index (34.02±1.19 vs. 33.72±1.35), failed to reach significance, Table 2.

**Table 2.** Summary of clinical outcomes using scar assessment tools. Scar characteristic assessed by qualitative questionnaire and quantitative measurements, including mVSS, DermaScan^®^ ultrasound, cutometer and DSMIII^®^ colorimeter.

| Scar Assessments | Category | Time points |  |  |  |
| --- | --- | --- | --- | --- | --- |
| Range,<br>(MEAN±SEM) | (n=14) | Baseline | M3 | M6 | Y1 (End point) |
| Modified Vancouver Scar Scale | Overall score | 3-7<br>(4.64±0.29) | 3-7<br>(5.14±0.27) | 3-7<br>(5.14±0.27) | 3-7<br>(5.14±0.31) |
|  | Pliability | 1-3<br>(1.64±0.20) | 1-3<br>(1.64±0.17) | 1-3<br>(1.71±0.19) | 1-3<br>(1.86±0.14) |
|  | Height | 0-1<br>(0.64±0.13) | 0-1<br>(0.71±0.13) | 0-1<br>(0.64±0.13) | 0-1<br>(0.71±0.13) |
|  | Vascularity | 0-1<br>(0.86±0.10) | 1-2<br>(1.07±0.07) | 1-1<br>(1.00±0.00) | 1-1<br>(1.00±0.00) |
|  | Pigmentation | 1-2<br>(1.50±0.14) | 1-2<br>(1.71±1.13) | 1-3<br>(1.8±0.15) | 1-2<br>(1.57±0.14) |
| DermaScan Ultrasound | Thickness | 1.75-3.93<br>(2.67±0.19) | 1.51-3.40<br>(2.45±0.18) | 1.80-4.75<br>(2.78±0.20) | 1.95-3.54<br>(2.64±0.12) |
| DSMIII Cutometer | Cutometer (R0) | 0.23-1.18<br>(0.66±0.08) | 0.17-1.11<br>(0.68±0.08) | 0.15-1.03<br>(0.66±0.08) | 0.19-1.13<br>(0.60±0.07) |
|  | Cutometer (R2) | 0.74-0.95<br>(0.86±0.02) | 0.70-0.96<br>(0.86±0.02) | 0.59-0.96<br>(0.84±0.03) | 0.69-0.96<br>(0.85±0.02) |
| Colourimeter | Erythema | 4.30-17.19<br>(9.27±0.95) | 4.56-15.74<br>(9.54±0.92) | 6.97-17.37<br>(10.69±0.86) | 6.32-16.73<br>(10.88±0.81) |
|  | Pigmentation | 31.81-41.77<br>(33.72±1.35) | 22.90-42.33<br>(33.02±1.80) | 29.19-41.21<br>(34.82±1.03) | 25.62-44.27<br>(34.02±1.19) |
|  | Chroma (Cab*) | 11.15-23.46<br>(16.36±1.11) | 11.92-23.07<br>(16.36±1.08) | 12.82-24.21<br>(17.58±0.98) | 8.29-38.51<br>(16.73±2.04) |
|  | L* (Lightness) | 26.68-64.74<br>(43.29±2.88) | 30.54-64.93<br>(44.11±3.02) | 32.27-62.42<br>(45.26±2.86) | 39.37-68.33<br>(59.49±2.43) |
|  | a* (Redness) | 7.18-19.23<br>(11.58±0.95) | 7.76-19.06<br>(12.30±0.84) | 8.79-17.61<br>(12.29±0.74) | 7.30-38.45<br>(13.47±2.05) |
|  | b* (Yellowness) | 4.98-19.99<br>(11.04±1.10) | 4.70-17.73<br>(10.46±1.00) | 7.05-21.45<br>(12.10±1.14) | 2.13-18.73<br>(8.39±1.46) |

As our participant’s scars varied greatly in how long they had been established, we considered this as a potential factor in laser efficacy. To investigate whether scar age was related to treatment outcomes, we first ranked scar outcomes from the best to worst response (rank 1 to 14) using the sum of the total scar assessment scores. This ranking revealed that participants with better overall scores for scar reduction had younger scars, Figure 1a. Therefore, to further identify differences in participants’ response to treatments, we divided them into those with younger scars, <6 years since wound closure (n=7) and those with older scars, > 6 years since wound closure (n=7). Demographics of these two groups are shown in Table 1. Our analysis showed that young scars had significantly better scar reduction than old scars in skin thickness (14.12±7.08% vs. −23.00±13.30%, p<0.05), mVSS overall score (−0.48±5.93% vs. −25.48±8.99%, p<0.05), and vascularity (−1.88±9.78% vs. −57.29±19.24%, p<0.05), Figure 1b-1d. There was also a trend towards better reduction in pigmentation for younger scars (4.64±3.41% vs. −58.98±5.82%, p=0.072), Figure 1e. AFCO_2_ laser treatment produced satisfactory colorimetric adjustment for all participants with no differences between young and old scars, Figure 1f-1i.

**Figure 1.**
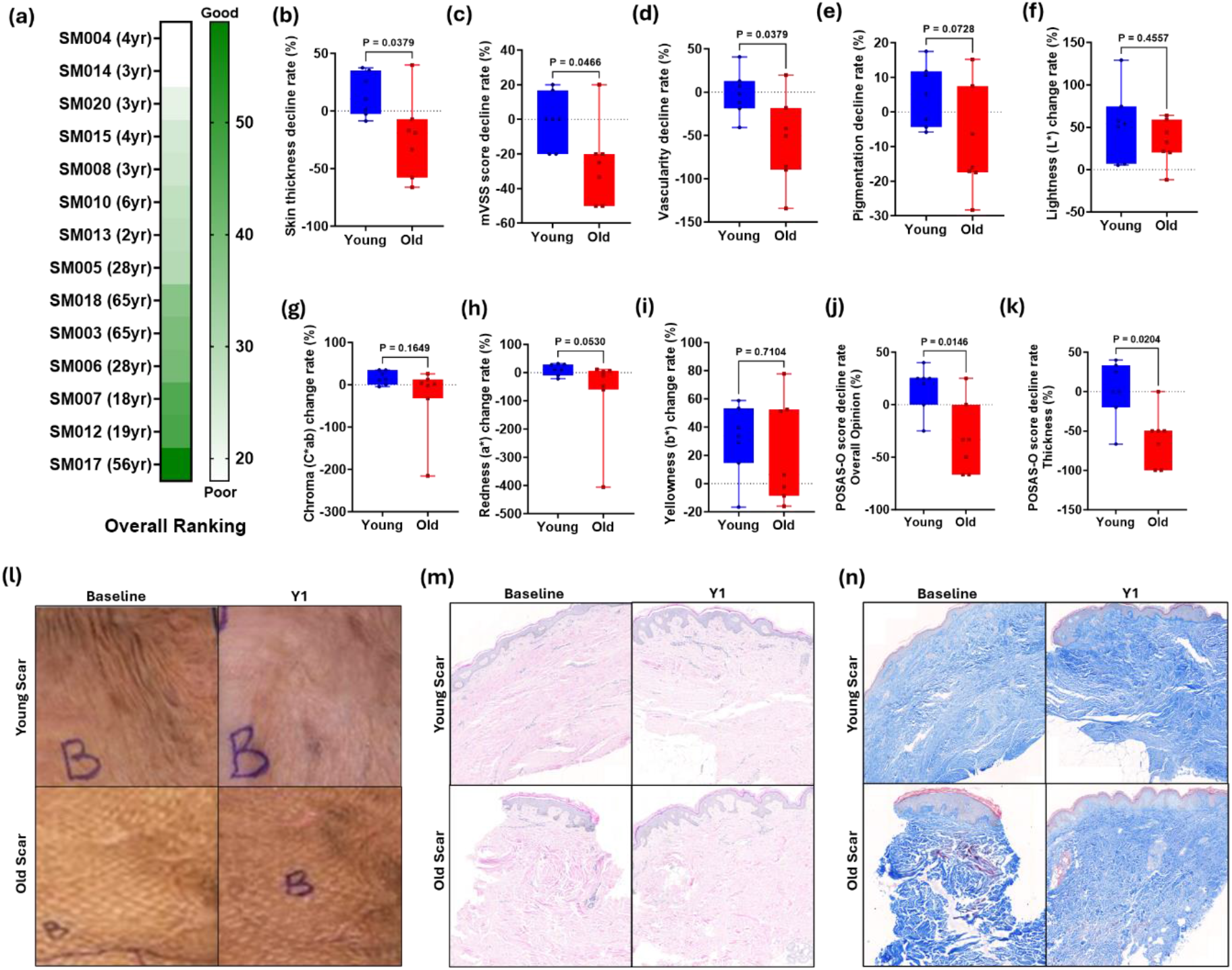
Laser treatment outcomes of young and old scars after AFCO_2_. (a) Overall ranking of scar reduction rate. Scar reduction rates were calculated using parameters including mVSS overall score, skin thickness measurements from Dermascan® ultrasound and data from DSMIII® colorimeter. Participants were ranked from the best to worst based on the scar reduction rates for each parameter (rank 1 to 14). Then, overall ranking is calculated based on the sum of the rankings for each individual parameters for each participant (good to poor, colour white to green). (b) Difference of scar decline rate between young and old scars in skin thickness, (c) mVSS overall score, (d) vascularity, (e) pigmentation, (f) lightness, (g) chroma *C*ab*, (h) redness, (i) yellowness, (j) POSAS-O overall score and (k) POSAS-O skin thickness. (l) Representative photographic images of young and old scars before and after AFCO_2_ treatment. (m) Representative H&E histological staining images of young and old scars before and after AFCO_2_ treatment. (n) Representative Masson Trichrome Blue staining images showing collagen arrangements of young and old scars before and after AFCO_2_ treatment.

These findings were further supported by POSAS-O, which also showed better treatment responses in younger scars compared to older scars (15.71±8.12% vs. -32.14±12.91%, p<0.05), and skin thickness (1.67±13.94% vs. -59.52±13.04%, p<0.05), Figure 1j-1k. Representative images recorded using Vectra 3D photography further confirmed that younger scars after treatment exhibited a smoother texture compared to older scars, Figure 1l. Finally, histological staining using H&E and Masson Trichrome blue staining, we found that the collagen arrangements in the young scars were more organised and structured than older scars post-therapy, Figure 1m-1n. From these clinical observations we concluded that overall, younger scars had better clinical outcomes than older scars following AFCO_2_ laser therapy.

### scRNAseq of skin biopsies reveals differences in cell content associated with differential scar reduction

Hypertrophic scars at different maturation stages may contain distinct populations that may determine responses to laser treatment. To explore the biological mechanisms potentially underpinning the differences in clinical scar outcomes post-laser therapy, single-cell RNA sequencing analysis (scRNAseq) was carried out on longitudinal full thickness skin biopsies obtained from two young scars (good responder, GR) and two old scars (poor responder, PR), at 4 time points (D0, W3, M6, Y1). Three healthy control (HC, mean age 54.67±6.49 years) samples were also analysed. In total, 116,759 cells were analysed, with an average of 20,431 reads per cell. Integrating the data from all samples and using unsupervised uniform manifold approximation and projection (UMAP), we identified nine major skin cell types based on established lineage-specific marker genes from all 3 groups (GR, PR and HC), these were: fibroblasts (*PDGFRA^+^*); vascular endothelial cells (VEC; *VWF^+^, SELE^+^, CLDN5^+^*); lymphatic endothelial cells (LEC; *LYVE1^+^*); pericytes (*ACTA2^+^, RGS5^+^, PDGFRB^+^*); keratinocytes (KRT; *KRT14^+^*); melanocytes (Mel; *MLANA^+^*); Schwann cells (Sch; *NRXN1^+^*); lymphoid cells (Lymphoid; PTPRC^+^, *CD3E^+^*) and myeloid cells (Myeloid; *PTPRC^+^, CD3E^-^, HLA-DRA^+^, CD68^+^*), Figure 2a-2d & Supplementary Figure 2a.

**Figure 2.**
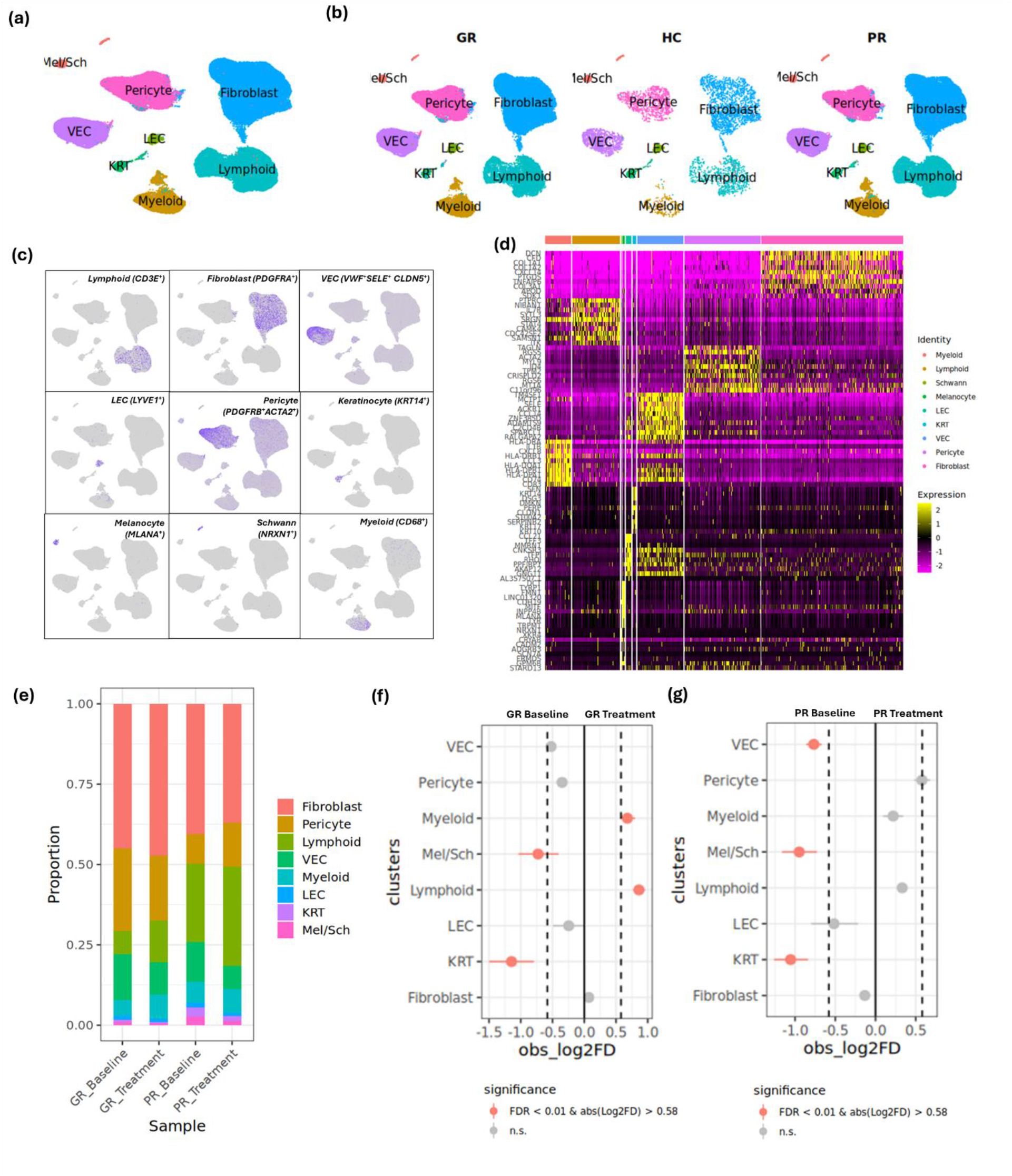
Single-cell RNA-sequencing reveals distinct cellular heterogeneity of skin scar tissues. (a) Unsupervised uniform manifold approximation and projection UMAP plot revealed cellular heterogeneity with 9 distinct clusters of skin cells identified and colour coded. (b) UMAP plots showing the distributions of skin cell clusters in healthy skin tissue (HS), young scar (GR) and old scar (PR) tissues. (c) Feature gene expression plots showing selected cluster-specific gene distributions in skin cells. (d) Heatmap showing representative differentially expressed genes in each identified cell types. (e) Cell proportions of skin cell populations in GR and PR tissues. Permutation test was used to test the significance in cell proportions between (f) GR baseline and GR treatment, (g) PR baseline and PR treatment.

Next, we analysed the pooled data for the proportions of each cell type in scar tissues, as well as healthy controls. We found that the proportions of fibroblast and immune cells was significantly higher in scars compared to healthy control biopsies, consistent with the pro-inflammatory and pro-fibrotic mechanistic nature of hypertrophic scar pathogenesis, Supplementary Figure 2b-2c. To take a closer look at the differences in cell proportional changes in response to laser therapy between GR and PR, we further compared scar samples at baseline (D0) for GR and PR, with pooled post-treatment samples for GR (D21, M6, Y1) and PR (D21, M6, Y1), Figure 2e. In GR, we found significant increases in immune cell populations (lymphoid and myeloid) after AFCO_2_, Figure 2f. In contrast in PR, although there was a slight increase in pericyte, lymphoid and myeloid cells, we didn’t find any significant changes in cell populations after AFCO_2_, Figure 2g.

### Good and poor response to laser therapy is associated with distinct gene expression

In order to further delineate the biological differences between GR and PR skin samples post AFCO_2_, we next studied their gene expression profiles. As shown in the hierarchical clustering heatmap analysed using Pseudobulk differential gene expression (DEG) software package in R, we found that gene expression profiles in skin samples differed significantly between groups, Supplementary Figure 2d. In order to take a closer look at the differences between GR and PR, we examined the DEG profiles and performed GO term enrichment analysis comparing the two outcomes. GO term enrichment analysis revealed that biological pathways including extracellular matrix organisation and collagen trimer formation were significantly enriched in both GR and PR after AFCO_2_ treatment. Pathways unique to GR scars after AFCO_2_ (GR Treatment) included extracellular matrix structure constituent and organisation, collagen fibril organisation and collagen binding, Figure 3a-3b. Genes, such as *COL14A1, POSTN, COL1A2, SPARC, KLK4, LUM and ADAMTS19,* can be further traced back to the fibroblast cluster, indicating that fibroblasts are one of the major players driving the good response to laser therapy (Figure 3c, 3e & 3f).

**Figure 3.**
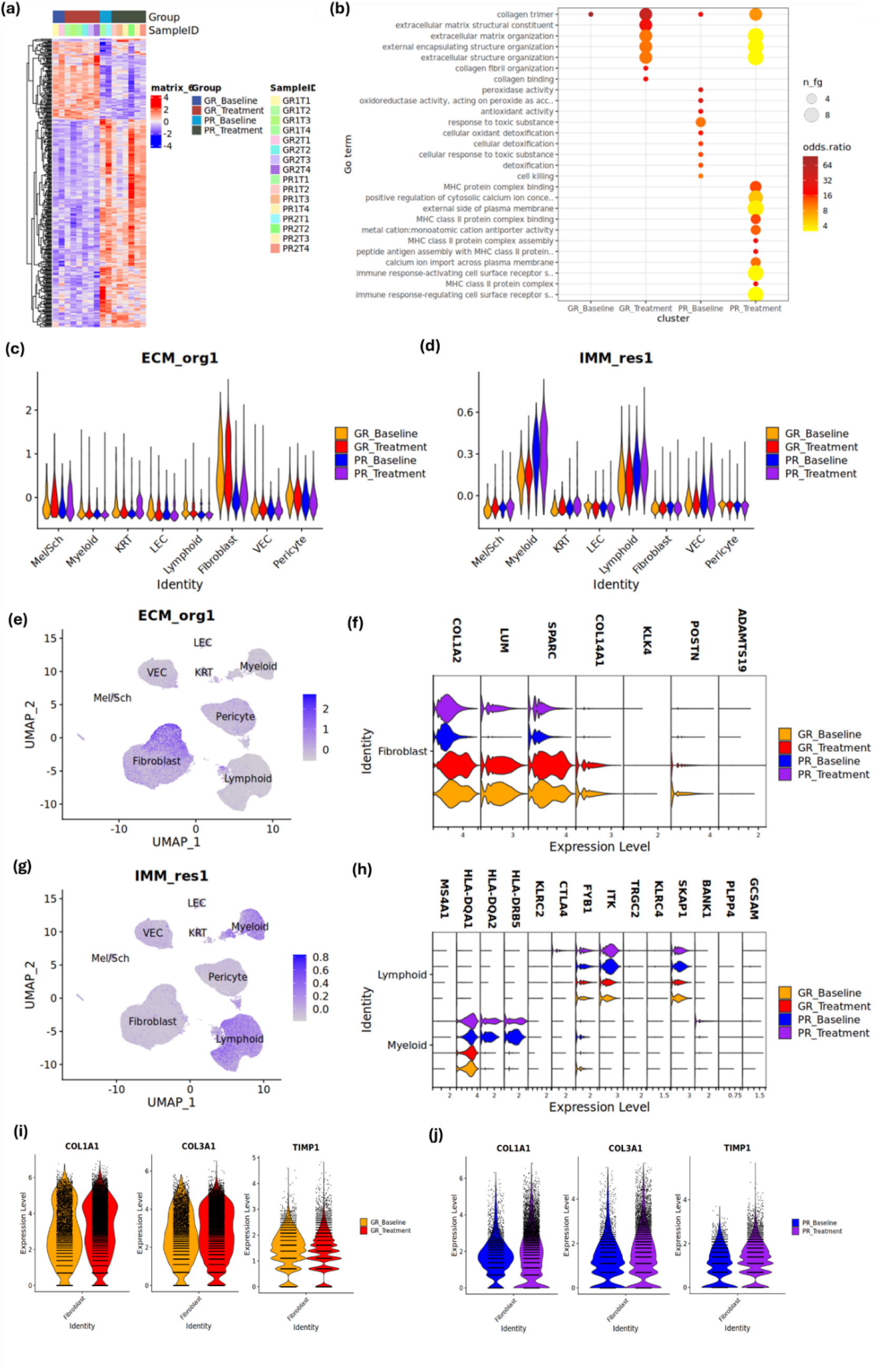
Pseudobulk differential expression analysis and functional analysis comparing normal and scar tissues. (a) Heatmap of differential gene expression in GR and PR tissues. (b) GO Term enrichment analysis of biological process, molecular function and cellular components of DEGs in skin cells between GR baseline, GR treatment, PR baseline and PR treatment. (c) Violin plot illustrating genes that were enriched in ECM organisations in GR were mapped to fibroblast population. (d) Violin plot showing genes that were enriched in immune responses in PR were mapped to myeloid and lymphoid cell populations. (e) Feature gene plot showing genes that were enriched in ECM organisations in GR were mapped to fibroblast population. (f) Violin plot illustrating the expression of some DEGs in fibroblast population between GR and PR. (g) Feature gene plot showing genes that were enriched in immune responses in PR were mapped to lymphoid and myeloid cell populations. (h) Violin plot illustrating the expression of some DEGs in myeloid and lymphoid populations between GR and PR. (i) Violin plots illustrating the expressions of *COL1A1, COL3A1 and TIMP1* in GR and PR scars before and after treatment. (j) Violin plots illustrating the expressions of COL1A1, COL3A1 and TIMP1 in GR and PR scars before and after treatment.

Biological pathways upregulated in PR scars post AFCO_2_ were associated with immune response-activating cell surface receptor signalling pathway and immune response-regulating cell surface receptor signalling pathway, Figure 3b. Despite no apparent increase in immune cell infiltration after laser therapy in PR (Figure 2g), genes such as *SKAP1, ITK, FYB1, HLA-DRB5, HLA-DQA2, HLA-DQA1, CTLA4, MS4A1, KLRC2, KLRC4, TRGC2, BANK1, PLPP4 and GCSAM*, can be traced back to lymphoid and myeloid cell populations, suggesting the activation of these immune cells as a crucial factor in driving poorer responses to laser treatment, Figure 3d, 3g & 3h.

As previous studies suggested that fractional CO2 laser treatments increase type III to type I collagen in mature burn scars^27, 28^, we further examined the expression of *COL3A1* and *COL1A1* in our scar biopsies. Furthermore, the expression of the pro-fibrotic factor, tissue inhibitors of metalloproteinase 1 (*TIMP1*)^29^, were also studied. We showed increased *COL3A1* and decreased *TIMP1* expression in GR after laser treatment, whereas the opposite was found in PR scars post treatment, Figure 3i-3j.

Collectively, these data suggest that fibroblasts and immune cells are key populations that may influence different AFCO_2_ laser treatment outcomes.

### Distinct gene expression patterns identified in fibroblast and immune cell populations from good and poor responders

Next, we investigated the gene expression profiles in cell sub-populations to further investigate the differences between GR and PR scars after AFCO_2_. We found that lymphoid cells, fibroblasts and to a lesser extent myeloid cells were the three major cell populations with the greatest differences in gene expression patterns post AFCO_2_ between GR and PR, Figure 4a-4c & Supplementary Figure 3. We found the largest number of DEG between GR and PR was observed in lymphoid cell populations (n=746, PR Treatment vs GR Treatment) and the main genes distinguishing the two groups include *FKBP5*, *RUNX2, PDE3B, SESN3*, *EPSTI1* and *LINC01619* in PR and *CCL4*, *PLCG2*, *S1004A, RPS29* and *XCL2* in GR, Figure 3i & Supplementary Figure 3. Fibroblasts are another cell population with highly variable gene expression (n=429, PR Treatment vs GR Treatment). Key genes that differentiate them in fibroblast populations including *APOD, PHLDA1, MT1X, RGS2, PLA2G2A, ID2, PTGS2, ADAMTS1, HSPA1B, ZFP36L2, JUN, GSN, KLF2, FOS, MGST1, GPX3, COL23A1 and TNC* in PR; and *COL1A1, COL3A1, SPARC, SDK1, CTHRC1, LUM, XIST, COL14A1, ASPN, POSTN, PDGFRL* and *PCOLCE* in GR, Figure 3j & Supplementary Figure 3.

**Figure 4.**
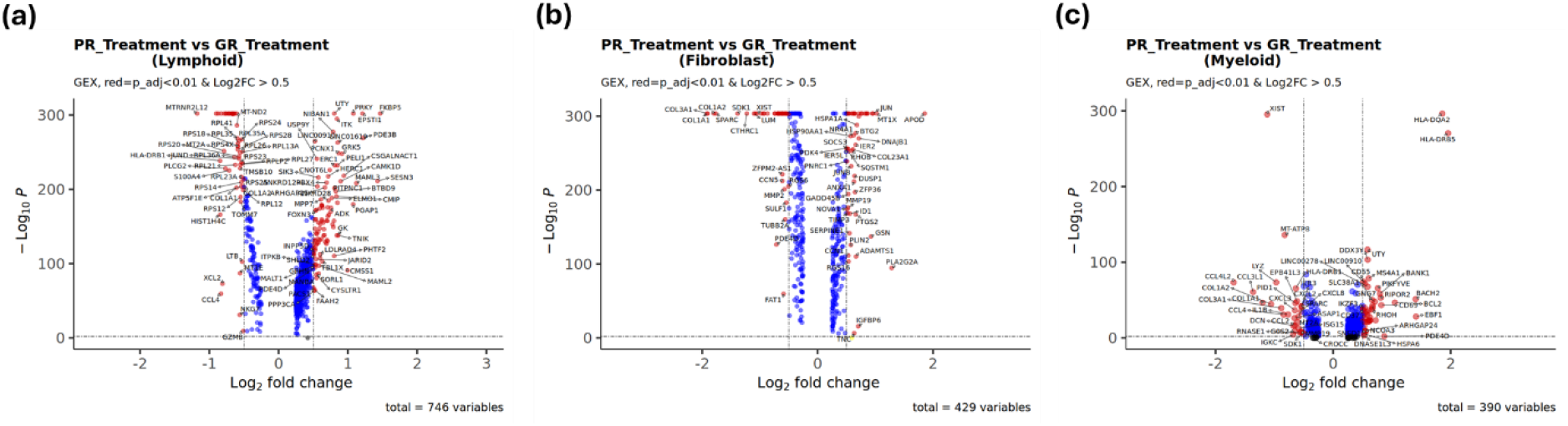
Distinct gene expression patterns identified between GR and PR post AFCO_2_. (a) Volcano plot of DEGs in lymphoid cell population between GR treatment and PR treatment. (b) Volcano plot of DEGs in fibroblast cell population between GR treatment and PR treatment. (c) Volcano plot of DEGs in myeloid cell population between GR treatment and PR treatment. GR1T1, Young scar 1, Baseline; GR1T2, Young scar 1, D21; GR1T3, Young scar 1, M6; GR1T4, Young scar 1, Endpoint (Y1); GR2T1, Young scar 2, Baseline; GR2T2, Young scar 2, D21; GR2T3, Young scar 2, M6; GR2T4, Young scar 1, Y1; PR1T1, Old scar 1, Baseline; PR1T2, Old scar 1, D21; PR1T3, Old scar 1, M6; PR1T4, Old scar 1, Endpoint (Y1); PR2T1, Old scar 2, Baseline; PR2T2, Old scar 2, D21; PR2T3, Old scar 2, M6; PR2T4, Old scar 1, Y1.

Pseudobulk analysis was then used to give cell type specific differences, Supplementary Figure 3. In fibroblasts, despite similar biological pathways in ECM organisation being observed in GR and PR, there were specific gene differences. Pathways associated with ECM structure constituent and collagen binding were specifically identified in GR Treatment and genes upregulated in these pathways included *SPARC, COL14A1 and CTHRC1*, Supplementary Figure 3a-3b. In addition, the pathways enriched in PR Treatment were also associated with myeloid leukocyte migration, as well as pathways relating to the regulation of nervous system development and musculoskeletal system responses, Supplementary Figure 3b. Of note, specific immune response related pathways were identified in PR Baseline prior to AFCO_2_, including interleukin-12 production, regulation of interleukin-12 production, positive regulation of interleukin-12 production and positive regulation of protein secretion, suggesting a potential role of immune response in differentiating GR and PR treatment outcomes, Supplementary Figure 3b. In lymphoid cells, genes enriched in PR Treatment were associated with regulation of leukocyte migration, negative regulation of myoblast differentiation and regulation of nervous system process. Conversely, the genes upregulated in GR were associated with humoral immune responses and positive regulation of T cell activation, Supplementary Figure 3d-3e. In myeloid cells, enriched pathways were identified only in PR after AFCO_2_. These pathways included, antigen processing and presentation of exogenous peptide antigen via MHC class II, immunoglobulin production, production of molecular mediator of immune response and adaptive immune response based on somatic recombination of immune receptors built from immunoglobulin superfamily domains, Supplementary Figure 3g-3h.

In summary, our data suggested that PR and GR scars have distinct cell population frequencies and patterns of gene expressions following laser therapy, with genes enriched in GR scars favouring ECM structural organisation, whereas genes enriched in PR scars favoured immune/inflammatory function. The increased expression of inflammatory gene signatures in PR scars hinted that these scars may remain in the inflammatory phase of the response to laser treatment without progressing to the later stage of tissue remodelling.

### Enhanced ECM intercellular communication networks between fibroblast and immune cells observed in PR scars with fibroblasts as the main orchestrator

As the wound healing process is a complex biological development which involves several cellular communication networks to support its progression, we wondered whether there were any specific networks that underpin the differences between GR and PR scar reduction after laser therapy. Therefore, we investigated the intercellular communication networks within the GR and PR scars. Using the CellChatDB algorithm to infer the intercellular communication networks within the ECM, we detected 73 enriched ligand-receptor genes that linked to 144 significant ligand-receptor pairs in GR Treatment (GRTx); and 78 enriched ligand-receptor genes that linked to 164 significant ligand-receptor pairs in PR Treatment (PRTx). These networks can be categorised in to 8 signalling pathways, including COLLAGEN, LAMININ, FN1, TENASCIN, THBS, VWF, HSPG and VTN pathways. Comparing GR and PR, we found that overall communication networks, such as the number of interactions and relative information flow, were increased in PR scars after AFCO_2_, Figure 5a-5b; with collagen, in particular, being the prominent pathway highlighting the differences, Figure 4a-4d. Using network centrality analysis to infer collagen signalling networks in both GR and PR, we identified that the fibroblast population is the most prominent source for collagen ligands acting onto myeloid cells. Fibroblasts were also the dominant mediator and influencer in the collagen network, suggesting a critical role in orchestrating cell-cell communication within the ECM network, Figure 4e-4f. Increased network traffic was also found in lymphoid cells which were identified as another important cell type that served as receivers in the PR collagen signalling network, Figure 4f.

**Figure 5.**
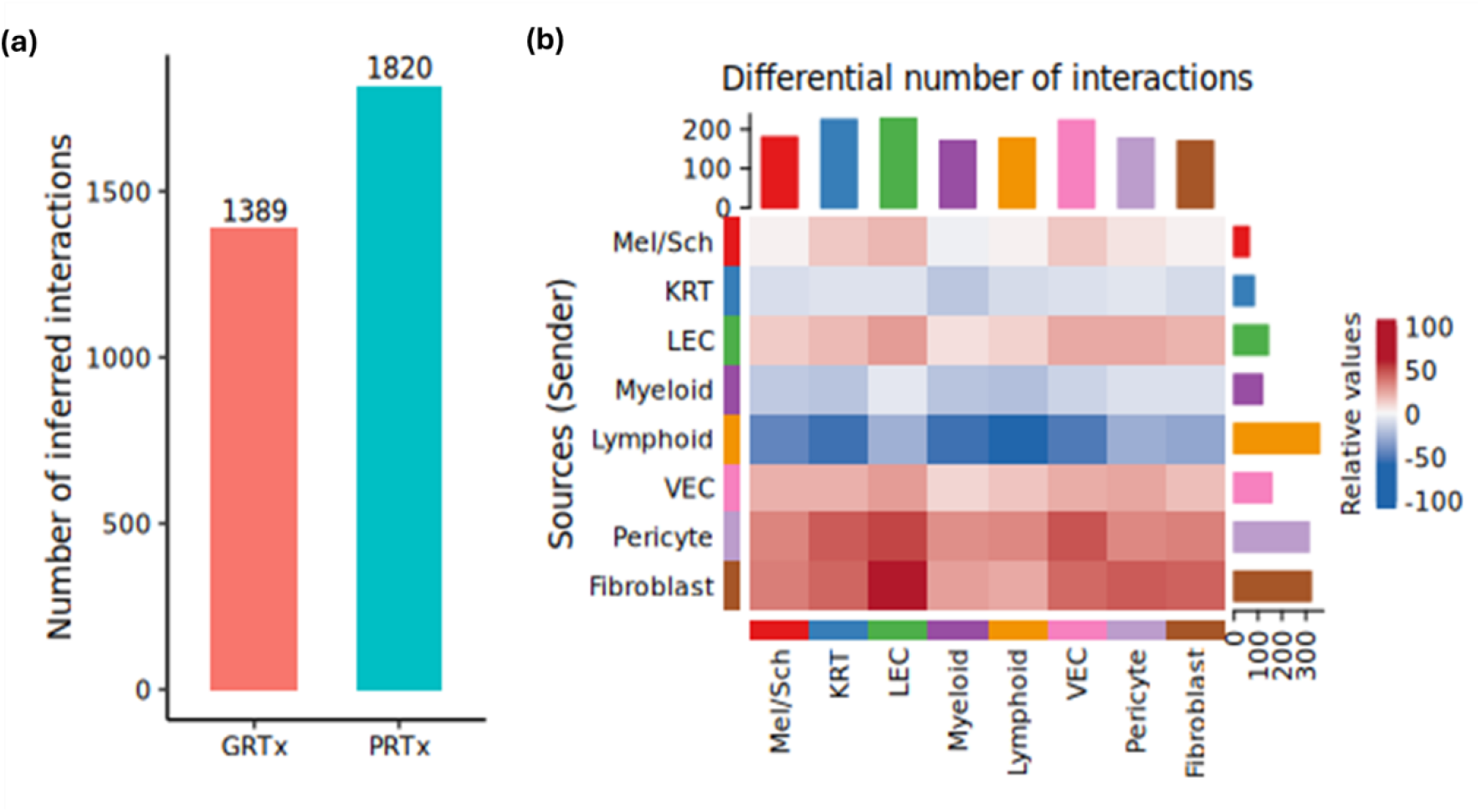
Increased ECM intercellular communication network found in PR scars after AFCO_2_. (a) Number of inferred interactions within ECM communications networks in GR treatment (GRTx) and PR treatment (PRTx). (b) Differential number of intercellular interactions in ECM between GRTx and PRTx. Red, increased number of ECM inter-cellular interactions in PRTx, compared to GRTx. Blue: decreased number of ECM intercellular interactions in PRTx, compared to GRTx.

**Figure 6.**
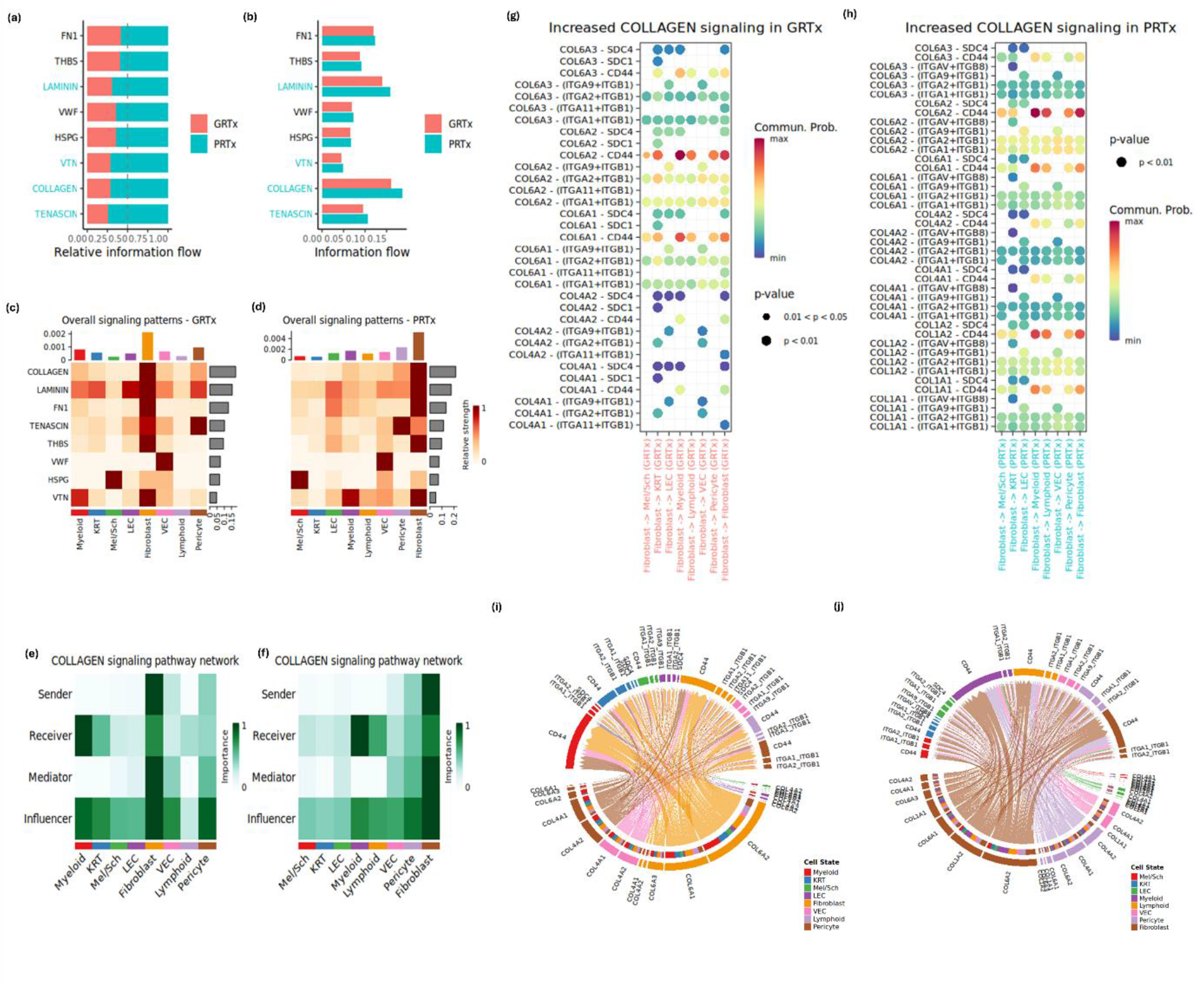
GR and PR scars have different patterns of ECM intercellular communication network, with fibroblast being the main driver in the network conversations. (a) Relative information flow of ECM pathways in GRTx and PRTx. Red, GRTx; Green, PRTx. ECM Pathways were ranked based on the overall information flow within the inferred networks between GRTx and PRTx. Significant pathways in PRTx were highlighted in Green on Y-axis. (b) Information flow of ECM pathways in GRTx and PRTx. Red, GRTx; Green, PRTx. Significant pathways in PRTx were highlighted in Green on Y-axis. (c) Overall ECM signalling patterns in GRTx. (d) Overall ECM signalling patterns in PRTx. (e) Collagen signalling pathways network in GRTx. (f) Collagen signalling pathways network in PRTx. (g) Ligand-receptor pairs of collagen networks in GRTx from fibroblast to target cells.. (h) Ligand-receptor pairs of collagen networks in PRTx from fibroblast to target cells. (i) Chord diagram of the collagen cellular communication network in GRTx. (j) Chord diagram of the collagen cellular communication network in PRTx.

This finding is consistent with our previous findings in DEG and GO enrichment analysis, from which we found that the PR population was enriched in pathways linking to regulation of immune cell migrations, Supplementary Figure 3e. Analysing collagen communication networks between fibroblasts and cells including myeloid, fibroblast, vascular endothelial cells and lymphoid cells, we further revealed that despite increased communication probability in COL6A2-CD44 and COL6A1-CD44 were observed in both groups, networks unique to PR were also identified, including COL1A2-CD44, COL1A1-CD44, COL1A2-(ITGA2+ITGB1) and COL1A1-(ITGA1+ITGB1), Figure 4g-4h& Supplementary 4a-4b. Fibroblasts were the main source of these signalling networks and they acted in autocrine and paracrine manners targeting cell populations including immune cells such as myeloid, lymphoid cells, as well as vascular endothelial cells, Figure 4i-4j & Supplementary Figure 4c-4h.

Taken together, these increased ECM communication networks between fibroblast and immune cells, again suggested a crucial role of these cells in the PR scar response to laser therapy.

### Enhanced profibrotic and immune cell activation network traffic observed in PR scars with myeloid and vascular endothelial cells being the primary communicators

Based on our data, we found that PR scars had increased inflammatory features. It is well established that the wound healing process is comprised of four major phases: homeostasis, inflammation, proliferation and tissue remodelling. During the inflammation phase, the site of injury sends out signals to recruit immune cells. In response, immune cells from the circulation transit across blood vessels and enter the microenvironment to exert their function. The increased immune activity we found in the PR indicates that communication between cells involved in this phase may also be increased. Therefore, in addition to ECM network, we also studied the secretion-signalling (SS) and cell-cell contact (CC) communication networks within GR and PR. In SS network, we detected 266 enriched ligand-receptor genes that linked to 299 significant ligand-receptor pairs in GR; and 276 enriched ligand-receptor genes that linked to 301 significant ligand-receptor pairs in PR, which can be further categorised in to 66 signalling pathways, including but not limited to CCL, CXCL, MIF, PDGF, PERIOSTIN, TWEAK and MSTN pathways. Similar to the ECM communication network, we found that overall SS communication networks were increased in PR scars, Supplementary Figure 5a-5b. Among them, CCL, CXCL and VISFATIN, were the major signalling pathways taking part in the SS communication network, with vascular endothelial cells (VEC) being the major player mediating the communications, Supplementary Figure 5c-5e.

Comparing the overall information flow of each signalling pathway, we found increased information flow in the VISFATIN signalling pathway in PR, including NAMPT-(ITGAS+ITGB1) and NAMPT-INSR, which contribute to increased communications between immune cells, fibroblast and VEC, were observed (data not shown). Furthermore, upregulation of the MSTN pathway after AFCO_2_ was found to be specific to PR, mediating communication traffic between endothelial cells, fibroblasts and myeloid cells, Supplementary Figure 5f.

In the CC network, we detected 180 enriched ligand-receptor genes that linked to 142 significant ligand-receptor pairs in GR; and 201 enriched ligand-receptor genes that linked to 158 significant ligand-receptor pairs in PR, which can be further categorised in to 52 signalling pathways. Again, we found that overall CC communication networks were also increased in poor responders, Supplementary Figure 5g-5h. Among them, MHC-II, SELE and MHC-I were the major signalling pathways taking part in the CC communication network, with myeloid cells being the major player in the CC communication, Supplementary Figure 5i-5k. Comparing the overall information flow of each signalling pathway, increased expression of CD80, L1CAM and VISTA were observed in PR and we further identified CD226 and CLEC pathway as significant to PR and CD39 as significant in GR, Supplementary Figure 5l.

In summary, these findings again indicated that GR and PR responses to laser therapy are biologically different, and they possess different intercellular communication patterns. We found increased communication traffic in both SS and CC networks in PR, with VEC and myeloid cells being the main communicators. Furthermore, this increased traffic was linked to profibrotic signalling and immune activation.

### Single cell trajectory analysis reveals distinct fibroblast developmental trajectories between GR and PR

Fibroblasts play a central role in the pathogenesis of hypertrophic scars and they exhibit high degrees of heterogeneity during the scarring process. Above, we have demonstrated that gene expression in the fibroblast population differed greatly between GR and PR, Figure 3d. Therefore, we decided to take a closer look at this population. First of all, we further divided fibroblasts into five subsets based on feature marker expressions^30, 31^, including secretory papillary fibroblasts (sPF; *PTGS2*^+^), secretory reticular fibroblasts (sRF; *MFAP5^+^PTGS2^+^*), pro-inflammatory fibroblast (InflamF; *APOD*^+^), myofibroblast (MyoF; *TAGLN^+^ACTA2^+^)* and mesenchymal fibroblasts (MscF; *MFAP5^+^ASPN^+^*), Figure 7a. Then, we compared the cellular composition of these fibroblast populations between GR and PR. We found that the proportion of the MscF subset was significantly higher in the GR fibroblast populations; on the other hand, sPF and InflamF populations were most prominent in PR, Figure 7b-7c.

**Figure 7.**
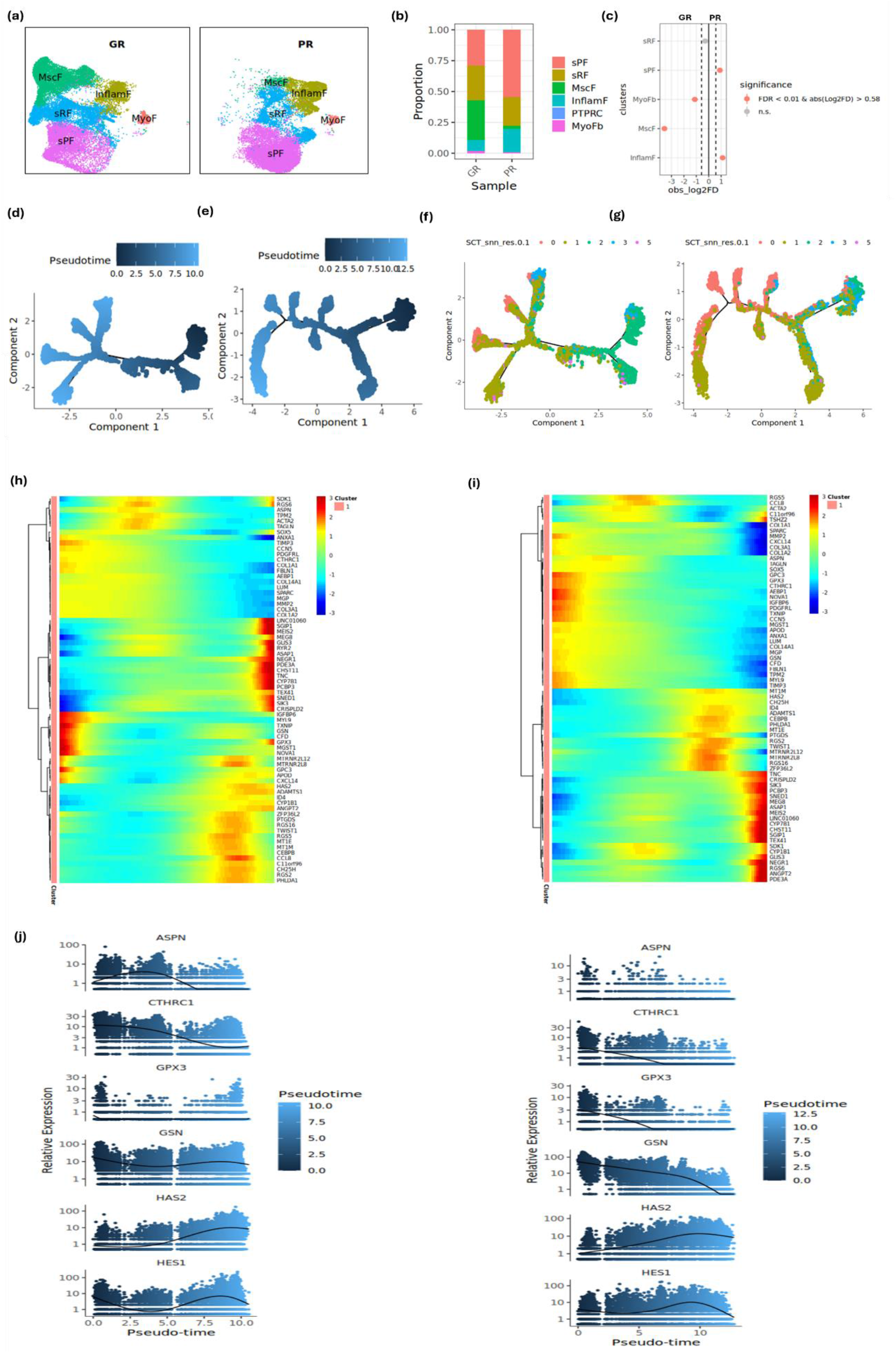
Pseudotime analysis reveals distinct fibroblast differentiation trajectories between GR and PR. (a) UMAP plots showing the distributions of fibroblast subpopulations in GR and PR. (b) Proportions of fibroblast subpopulations in GR and PR. (c) Permutation tests were used to test the significance in proportions of fibroblast subpopulations between GR (left) and PR (right) (Permutation test, FDR<0.01, abs. (Log2FD)>0.58, red dot). (d) Potential developmental trajectory of fibroblasts in GR inferred using *Monocle 2* software. (e) Potential developmental trajectory of fibroblasts in PR. (f) Mapping of GR’s fibroblast subpopulations in developmental trajectory. 0, sPF; 1, sRF; 2, MscF; 3, InflamF; 5, MyoF. (g) Mapping of PR’s fibroblast subpopulations in developmental trajectory. 0, sPF; 1, sRF; 2, MscF; 3, InflamF; 5, MyoF. (h) Heatmap visualisation of pseudotemporal expression pattern in GR. (i) Heatmap visualisation of pseudotemporal expression pattern in PR. Comparing relative gene expression patterns across pseudotime between GR and PR by (j) pseudo-time, (k) fibroblast subpopulations, (l) groups.

Finally, we performed single cell trajectory analysis using *Monocle 2* software package on fibroblast populations to study the developmental state overtime. By ordering fibroblasts into a major trajectory, we found that GR and PR undergo distinct developmental trajectories, with 9 and 11 developmental states, respectively, Figure 7d-7g. Interestingly, mapping fibroblast subsets along the developmental trajectories, we found an abundance of MscF presented at early stages of the GR development trajectory. Gene changes across pseudotime in GR and PR were further visualised using *pheatmap* software package. Since both groups are essentially hypertrophic scars, overall, they shared similar gene expression patterns across pseudotime, Figure 7h-7i. Despite this, varying differential gene expression patterns were also observed, Figure 7j. *ASPN* was highly expressed in GR but absent in PR. Its expression was found to increase at early stages of laser treatment, which then declined overtime. Although *CTHRC1* was expressed in both GR and PR, its expression was higher in GR at early stages of the laser response. On the other hand, *GPX3* expression also appeared at the early stage but its expression was significantly higher in PR. Likewise, *GSN* expression was also significantly higher in PR, but its expression declined sharply at later stage of the response whilst GR still maintained certain levels of expression. *HAS2* and *HES1* were both expressed GR and PR, however, the expression levels were absent in GR at the early stage.

Taken together, we showed GR and PR have distinct proportions of fibroblast subpopulations, with GR comprised of higher levels of regenerative MscF and PR comprised of higher levels of inflammatory InflamF and sPF populations. GR and PR underwent different developmental trajectories during the response to laser treatment. Higher levels of regenerative MscF from GR mapped to earlier stages of the response, together with distinct gene expression patterns across time may explain the reason driving the differences.

## Discussion

To summarise, our study demonstrated that AFCO_2_ laser treatment is more effective on less well-established scars, with better scar reduction as assessed by skin thickness, vascularity, and mVSS overall scores. Our data showed that young scars and old scars showed different gene expression patterns in response to AFCO_2_ laser treatment, suggesting a relationship with the response to laser therapy. Genes enriched in young scars and thus a positive response, were associated with extracellular matrix and structure organisation; whereas genes enriched in old scars and associated with a poorer response were related to enhanced immune and inflammatory responses. Intercellular communication networks were enhanced in poor responders, with increased crosstalk between fibroblasts, immune cells and vascular endothelial cells. Moreover, different proportions of fibroblast subpopulations were observed with differential responses to laser therapy, specifically a good response was associated with higher proportions of regenerative MscF in the treated skin, whereas a poor response was associated with higher proportions of inflammatory InflamF and sPF. Single cell trajectory analysis further revealed that good and poor responders underwent different trajectories across time; the genes involved in the good response developmental trajectory were associated with regenerative features while in poor responders’ genes were associated with excessive fibrosis. These differences collectively appear to determine the fate of the scar response to AFCO_2_ laser treatment and ultimately influence treatment outcomes.

Clinically, the optimum timing for treating hypertrophic burn scars using AFCO_2_ laser therapy remains unclear. In recent years, there have been a few studies investigating the efficacy of early laser treatment on hypertrophic burn scars. For example, in a retrospective review Tan et al assessed the potential of early intervention on improving HTS using AFCO_2_. They analysed 223 HTS patients treated by AFCO_2_ laser within 12 months of injury and concluded the early AFCO_2_ treatment is safe and effective^32^. A recent randomised control trial (RCT) was conducted in USA to investigate the use of AFCO_2_ laser treatment on acute traumatic injuries to prevent scar formation and they showed that early AFCO_2_ laser treatment in acute traumatic wounds 3 months after injury resulted in significant reduction of scar formation^33^. Moreover, another recent RCT trial carried out in Western Australia investigated the impact of AFCO_2_ laser on patient-reported outcome measures (POSAS), subjective scar appearance, dermal architecture, and gene transcription in early burn scars. It concluded that AFCO_2_ laser treated scars significantly altered scar thickness and texture and presented better appearance after treatment. They also reported significant decrease in thick collagen fibres and significant increase in finer collagen fibre density post AFCO_2_. Using bulk RNAseq analysis, they further suggested that AFCO_2_ induces sustained changes in fibroblast gene expression, indicating potential beneficial mechanisms of AFCO_2_ in HTS^34^. Similar to the above-mentioned studies, our results showed early laser treatment gave better scar reduction in skin thickness, vascularity, and mVSS overall scores.

Laser therapy is commonly used in post-burn scar clinics and has been shown to relieve scar tension, breakdown local scar matrix and mobilise fibroblasts to generate new collagen for better tissue remodelling outcomes^9, 14–18^. Laser therapy essentially creates a new wound and attempts to induce a less fibrotic response than the original injury but the cell and molecular factors influencing laser efficacy remain poorly understood. Using a transgenic mouse model, Phan et al identified the *Wnt* effector transcription factor *Lef1* as a regenerative factor in fibroblasts of developing skin, that transforms adult skin to regenerate rather than scarring^35^. In contrast, Mascharak et al., showed that postnatal *Engrailed-1* (*En1*) activation in fibroblasts leads to excessive fibrosis and scarring^25^. Recent studies on keratinocyte dedifferentiation pathways provide further evidence on the involvement of forkhead box O1/Krüppel-like factor 4 (*Foxo1/Klf4*) pathway with skin regeneration after mechanical stretch and identified that E-cadherin (*Cdh1*) as a crucial factor for skin regeneration^36, 37^. To find out whether GR scars also exhibited similar regenerative features, we analysed specific gene expression, including *LEF1, TRPS1, CDH1, FOXO1* and *KLF4* in GR and PR scars. We showed that the *LEF1* gene expression level increased in both GR and PR scars after AFCO_2_, whereas *TRPS1, CDH1, FOXO1* and *KLF4* were only increased in GR scars after treatment and these same genes were reduced in PR after AFCO_2_, further indicating the transcriptomic differences between good and poor responders.

Aberrant inflammation during wound healing is associated with impaired healing and pathological scarring^38, 39^. In our study, we showed significantly higher proportions of immune cells in PR compared to GR biopsies at baseline. We also found differences between GR and PR in the baseline samples. In fibroblasts, specific immune response related pathways were identified in PR Baseline prior to AFCO_2_, including interleukin-12 production, regulation of interleukin-12 production, positive regulation of interleukin-12 production and positive regulation of protein secretion. Previous studies have shown that IL-12 is anti-angiogenic^40^ and serves as a chemoattractant for macrophages^41^. Using IL-12/IL-23 knockout mice, Matias *et al*., further revealed that IL-12 and IL-23 influence wound healing by modulating early inflammatory responses and subsequent angiogenesis^42^. Therefore, the observation of IL-12 in PR baseline indicates an underlying inflammatory predisposition prior to AFCO_2_, which may underlie the increased myeloid cell migration after treatment. In addition in lymphoid cells, genes enriched in PR after laser treatment were associated with immune activation, e.g. regulation of leukocyte migration, negative regulation of myoblast differentiation and regulation of nervous system process. In myeloid cells, enriched pathways identified in PR after laser treatment included pathways associated with cell activation, including antigen processing and presentation via MHC-II, as well as mediators of immune responses. In addition, studying DEG profiles between GR and PR scars, we also noticed increased expression of *HLA-DQA1*, *HLA-DQA2* in immune cell populations in PR scars, which may be other potential factors that modulate the increased inflammatory responses in PR scars. To mitigate these prolonged inflammations, potential multi-modal treatments or the use of steroids could help to control excessive inflammation and improve therapeutic outcomes.

Fibroblasts are the main cell population in the skin dermis. Our data suggest that in young and old scars, the composition of fibroblast subpopulations induced by the laser treatment is different, with young scars (good responder) comprised higher proportions of regenerative MscF which would promote new skin formation^43^, and old scars (poor responder) comprised of higher proportions of inflammatory InflamF and sPF which would be more conducive to fibrosis and thus prevention of scar reduction. Studies have indicated that inflammatory fibroblasts produce inflammatory cytokines and may be involved in inflammation progression and in the development of other pathogenic conditions, such as pain^44, 45^. Our data echoes these findings, and we showed that pathways enriched in PR after laser treatment were also associated with myeloid leukocyte migration, as well as pathways relating to the regulation of nervous system development and musculoskeletal system responses. Further clinical investigations into neurological changes, such as pain would provide more information in this area.

The increased proportion of InflamF in PR suggests a potential prolonged inflammation phase after AFCO_2_. Because fibroblast subsets are responsible for different biological functions^46^, the stimulation of fibroblasts in young and old scars by AFCO_2_ treatment could therefore lead to alternative outcomes. Comparing differential gene expression profiles and intercellular communication patterns between good and poor responders, we found poor responder biopsies were enriched with pro-inflammatory features and have enhanced cross-talk between fibroblasts and immune cells. Moreover, we found good responders and poor responders had different gene expression features that are responsible for distinct biological functions. Genes enriched in GR were prone to ECM structure and organisation and genes enriched in PR were associated with enhanced immune responses. Additionally, we found that in GR scar, the expression of *COL3A1*, which is commonly found in foetal skin, increased after laser treatment; and the expression of the pro-fibrotic factor, tissue inhibitors of metalloproteinase 1 (*Timp1*), decreased after laser treatment; whereas the opposite was found in PR scars post treatment (Supplementary 2d-2i). Adding to this phenomenon, inferring enhanced ECM intercellular communication, we found PR had enhanced intercellular communications between fibroblast and cells including myeloid, lymphoid, vascular endothelial cell and fibroblasts, in which they used ligand – receptor pairs unique to old scars, COL1A2-CD44, COL1A2-(ITGA2+ITGB1) and COL1A1-(ITGA1+ITGB1) for the communications.

Finally, inferring intercellular communication networks, we found that regulators of wound healing including *POSTN, TWEAK* and *PTN* were significantly elevated in GR. Despite periostin^+^ fibroblast was often associated with pro-fibrotic features^30^, it has also been previously reported to have several positive functions in scar formation, including promoting proliferation and differentiation of fibroblasts, re-epithelialisation and wound closure^47^. Others have reported that periostin can promote tissue repair by enhancing the function of adipose-derived stem cells^48^. TNF-related weak inducer of apoptosis (*TWEAK*) is another factor that exhibits beneficial roles in scarring. It has been shown to facilitate angiogenesis and regulate the production of mediators that are crucial for tissue regeneration^49^. In contrast, signalling associated with profibrotic and T cell activation pathways, including CD226^50^, MSTN^51^ and IL16^52^, were significantly increased in PR. Many studies have described the involvement of MSTN in excessive scarring and it has been indicated as potential target for skin healing^51^. For example, a study using full thickness burns rodent model found a 4-fold increase in MSTN^53^. Other studies using MSTN-*null* mice showed higher tissue regeneration and reduced fibrosis^54, 55^. Therefore, to improve PR outcomes, drugs targeting MSTN pathway could also be used as a supportive treatment, in addition to laser therapy.

There are some limitations to our study. For example, the number of samples in this study is relatively small. Therefore, further studies using a bigger sample size may be required to further understand the biological mechanisms involved in scar reduction after AFCO_2_. Moreover, recent studies have now suggested an association between the early laser therapy (i.e. within 12 months of injury) and improved treatment outcomes^32–34^, indicating the potential benefit of initiating laser treatment early for post-burn scars. The age of young scars in our study was < 6 years with 2 years being the youngest scar. Therefore, further investigating the properties of ultra young scars (<1 years) may provide us with further insight into the optimum time for treating scars with laser therapy.

To conclude, our study demonstrated that young scars and old scars are biologically distinct in their response to AFCO_2_ laser treatment. The differences in gene expression profiles and unique intercellular communication networks between them ultimately determine the fate for different treatment outcomes. Our findings suggest that young scars favour wound healing and regeneration, whereas old scars are prone to maintained inflammation and fibrosis, which potentially limits the ability to reduce scars with AFCO_2_. Therefore, better scar reduction may be seen if AFCO_2_ laser therapy is applied sooner after wound healing.

## Methods

### Study participants and study schedule

Adults with established hypertrophic scars following a burn injury were recruited to the study. Participants gave their written informed consent and the study was approved by a UK National Research Ethics Committee, REC ref. 19/NS/0125). They received 3 sessions of AFCO_2_ laser treatments, at 3 month intervals over a 12-month period. During the study various validated scar assessment procedures were used to evaluate scar characteristics before each treatment and at the endpoint (baseline, M3, M6, Y1). Skin biopsy samples were collected from participants before treatment (baseline), and 3 weeks, 6 months and 1 year after the 1^st^ laser treatment^26^.

### AFCO_2_ settings

The Lumenis® UltraPulse® CO2 laser device with DeepFX^TM^ and/or SCAARFX^TM^ headpiece were used. The treatment included a single pass of the following settings: (1) SCAARFX™: Energy: 110-150mJ, density: 3%, shape: 2, size: 10, pulses 1; and (2) DeepFX ^TM^: Energy: 17.5-20.0mJ, density 5%, shape: 2, size: 10, pulses: 1. Repeat rate 0.5 seconds; Frequency 300Hz; no active cooling was used during the treatment.

### Clinical Scar assessment measures

Clinical scar parameters, such as skin thickness, pliability, vascularity and pigmentation, were obtained from participants before each laser treatment and at the 12 month endpoint to evaluate treatment outcomes. Assessments used to collect scar parameters including modified Vancouver Scar Scale (mVSS), Dermascan® ultrasound, Cutometer and DSMIII® Colorimeter. In addition, Vectra 3D photography and POSAS-O were used as auxiliary measures.

### Modified Vancouver Scar Scale (mVSS)

An adapted version of the modified Vancouver Scar Scale (mVSS) was used to evaluate the clinical efficacy of the laser treatment^56–58^. The mVSS is a numerical assessment for skin characteristics including pliability (range, 0-5), height (range, 0-3), vascularity (range, 0-3) and pigmentation (range, 0-3). The mVSS of each patient was individually assessed by an experienced observer who was blinded to the study. The accessor scores each characteristic based on the difference between the site of evaluation and the normal skin.

### Dermascan® high frequency ultrasound

Dermascan C® USB (Cortex Technology ApS, Denmark)^59^ provides 20mHz B-scanning at 60 x 150 micron resolution, with 13mm penetration^60^, which allows the visualisation of soft tissue at high resolution providing a measurement of the skin thickness possible. Skin thickness is defined as the distance between the stratum corneum and the inner surface of the burn scar in the dermis.

### Cutometer® skin elasticity meter

The Cutometer® skin elasticity meter (MPA 580, Courage and Khazaka GmbH, Germany)^56, 61^ is used to evaluate the skin elasticity parameters, measuring the vertical deformation of skin in millimetres (mm) when pulled by a controlled vacuum into a circular aperture. Skin elasticity is expressed as absolute parameters (Ua, Ue, Uf, Ur and Uv) or relative parameters (R-, F- and Q-parameters)^62, 63^. R0 and R2, were used in this study: R0 (=Uf) was used to describe the stiffness (pliability/firmness) of the skin by measuring the maximum of kin deformation. R2 (=Ua/Uf) was used to describe visco-elasticity of the skin by measuring the ratio of final retraction and maximum deformation.

### DSMIII Colorimeter®

DSMIII colorimeter measures the colour feature of the skin^64^. It combines two colour system readings into one measurement: narrow-band spectrophotometry (melanin and erythema) and tristimulus reflectance colorimetry (CIE L*a*b* system, CIELAB), which allows us to quantitatively measure the skin vascularity (erythema index), pigmentation (melanin index) and colour shift during the AFCO_2_ therapy over time.

All colorimetric measurements are presented using CIELAB format in 3D space: lightness (L*= 0, black; to L* = 100, white), redness (a*) and yellowness (b*). The chroma (C*ab) represents the length in (a*, b*) space: 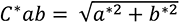. The CIELAB colour difference metric delta E (ΔE) between baseline and endpoint was calculated to quantify the overall colour shift: 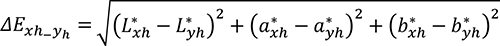 where *xh* correspond to endpoint (Y1) and *yh* correspond to baseline (D0).

### Patient Observer Scar Assessment Scale - Observer (POSAS-O)

The POSAS-O measures the quality of the scars by evaluating six dimensions including: Pain, itchiness, pliability, thickness, colour and irregularity from observers’ perspectives. Each dimension was scored from scale 1 to 10 based on the differences between the site of evaluation and the normal skin, with 10 indicating the largest difference. Additionally, the observer records their overall opinion score, from 1 to 10, regardless of the total score across the six evaluation dimensions.

### Human skin biopsy samples

5mm skin punch biopsy samples were obtained from participants undergoing AFCO_2_ laser treatment (baseline, week 3, month 6, and Y1). All biopsies collected were processed immediately and bisected with one half transferred to 10% neutral buffered formalin for downstream histological and immunohistochemical assessments; and the other half was fresh frozen in CS10 CryoStor® cell cryopreservation media at -80°C for future analysis using single cell RNA sequencing (scRNAseq) to determine cellular heterogeneity and transcriptomic differences in the scars.

### scRNAseq analysis

Skin biopsy samples (n=16) were thawed and washed with Dulbecco’s Phosphate-Buffered Saline (DPBS) supplemented with 5% foetal calf serum (FCS) once before incubating with 1mg/ml collagenase P (Sigma Aldrich, UK) and 100μg/ml DNase I (Sigma Aldrich, UK) for 3.5 hours at 37°C in a humidified CO2 incubator. Cell suspensions were filtered through a 70μm cell strainer. Dissociated cells were then labelled with live and dead dye before live cell sorting using a BD LSR FORTESSA™ X-20 Cell Analyzer (BD Biosciences, USA). Single cells were captured and barcoded using 10x Chromium platform and the sequencing libraries were prepared following manufacturer’s instructions using dual index Chromium Next GEM G Single Cell 3’ kit v3.1 (10xGemonics, California, USA). Libraries were quantified using an Agilent Bioanalyzer High Sensitivity DNA chip and pooled together to get similar numbers of reads from each single cell before sequencing. Overall, approximately 50,000 cells per sample were used for construction libraries and sequenced using Novogene single cell RNA sequencing services. Using Cell Ranger pipeline v.3.1.0 (10xGenomics), raw sequencing data were aligned to the hg38 reference genome and converted into a cell-feature matrix. This was followed by the unique molecular identifier (UMI) counting. In addition to 16 skin scar samples, 3 healthy control skin sample (age 44-67 years) sequencing data were integrated into our database for downstream analysis. We filtered out cells that had unique feature gene counts of less than 200 and cells that were contaminated with more than 15% mitochondrial counts. Transformed data were processed and refined before further data visualisation and differential analysis using, Seurat (Version 4.3.1)^65^ and *Monocle 2* (version 2.18.0)^66^ R packages and CellChatDB (v.3.0)^67^. Additionally, Loupe browser (10xGenomics) was used for data inspection and data visualisation. Full details of the analysis can be found in the online data supplement.

## Statistical analysis

Non-parametric Mann Whitney *U* test, Wilcoxon matched pairs signed rank test, Friedman test with multiple comparisons and One-Way ANOVA Statistical analysis with post-hoc test were performed using GraphPad Prism (v10.1.0) and R software.

## Acknowledgements

This study was funded by the Scar free Foundation. JML is supported by the NIHR Surgical Reconstruction and Microbiology Research centre, the views expressed here are those of the authors and not necessarily those of the NHS, NIHR or the Department for Health and Social Care.

---

**Supplementary Figure S1.**
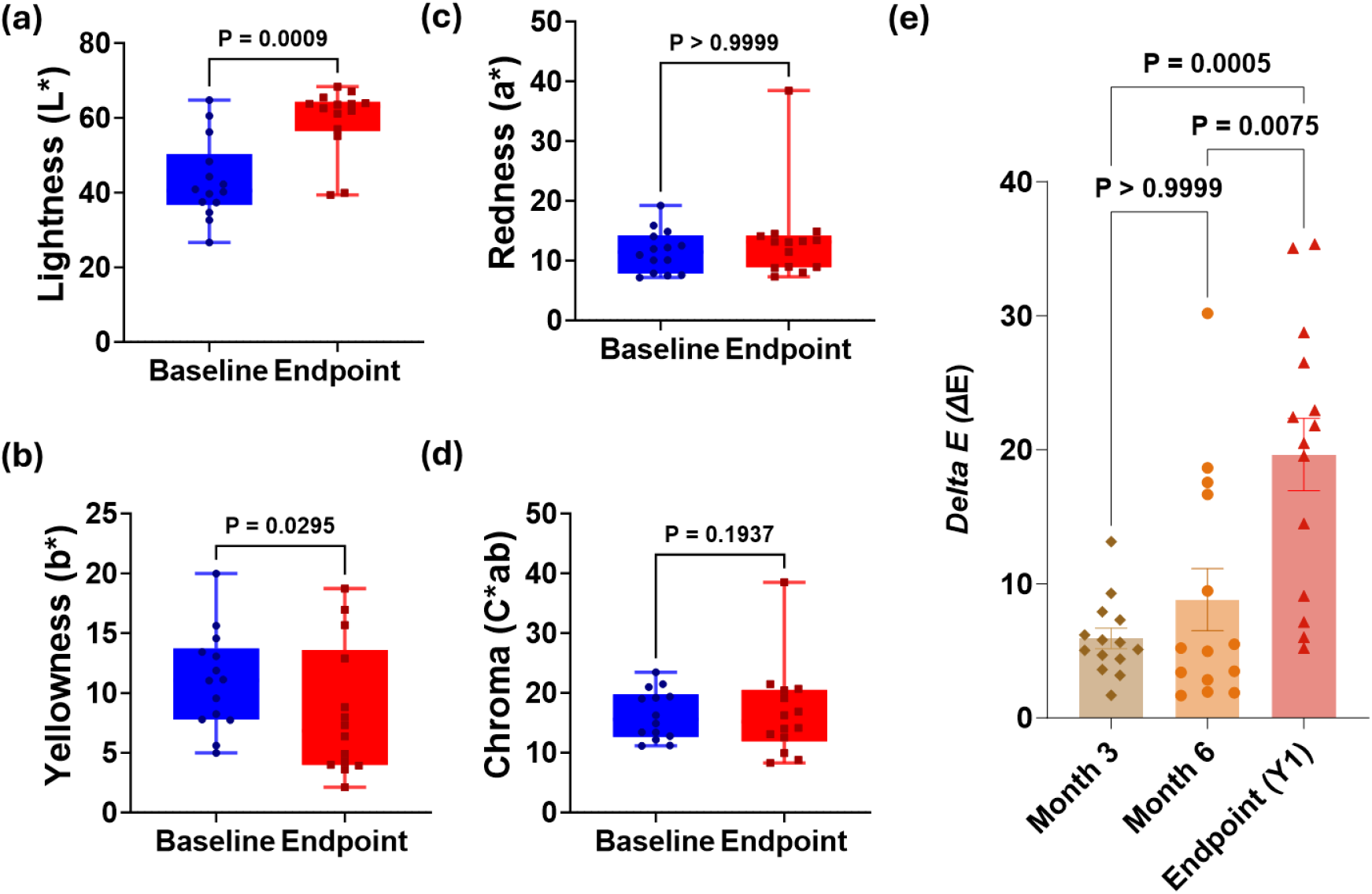
Colorimetric properties of scars. Differences in skin colour distribution between baseline and endpoint in (a) lightness, L*; (b) yellowness, b*; (c) redness, a* and (d) chroma, *C*ab*. Data presented as MEAN±SEM. Wilcoxon matched pairs signed rank test. (e) Skin colour difference (ΔE) from baseline to month 3 (Month 3), month 6 (Month 6) and endpoint (Y1). One Way ANOVA, Friedman test with Dunn’s multiple comparisons.

**Supplementary Figure S2.**
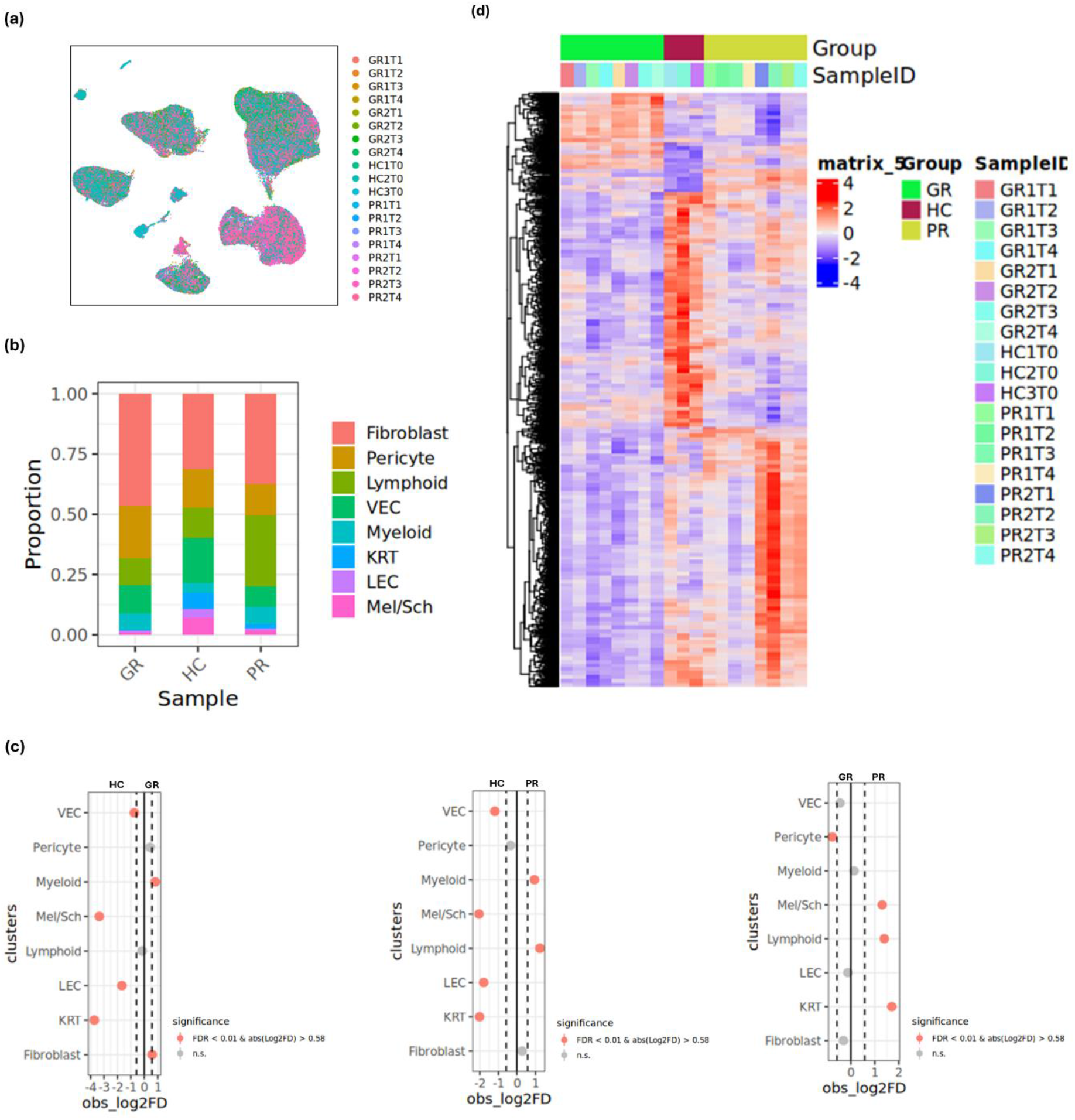
Pseudobulk differential expression analysis and functional analysis of skin cell subpopulations. (a) UMAP plots showing colour-coded skin samples mapped onto skin cell clusters. (b) Cell proportions of skin cell populations in HC, GR and PR tissues. (c) Permutation test was used to test the significance in cell proportions between HC and GR, HC and PR, as well as PR and GR (Left to right, FDR<0.01, abs. (Log2FD)>0.58, Permutation test, red dot). (d) Heatmap of differential gene expression between HS, GR and PR tissues. GR1T1, Young scar 1, Baseline; GR1T2, Young scar 1, D21; GR1T3, Young scar 1, M6; GR1T4, Young scar 1, Endpoint (Y1); GR2T1, Young scar 2, Baseline; GR2T2, Young scar 2, D21; GR2T3, Young scar 2, M6; GR2T4, Young scar 1, Y1; PR1T1, Old scar 1, Baseline; PR1T2, Old scar 1, D21; PR1T3, Old scar 1, M6; PR1T4, Old scar 1, Endpoint (Y1); PR2T1, Old scar 2, Baseline; PR2T2, Old scar 2, D21; PR2T3, Old scar 2, M6; PR2T4, Old scar 1, Y1.

**Supplementary Figure 3.**
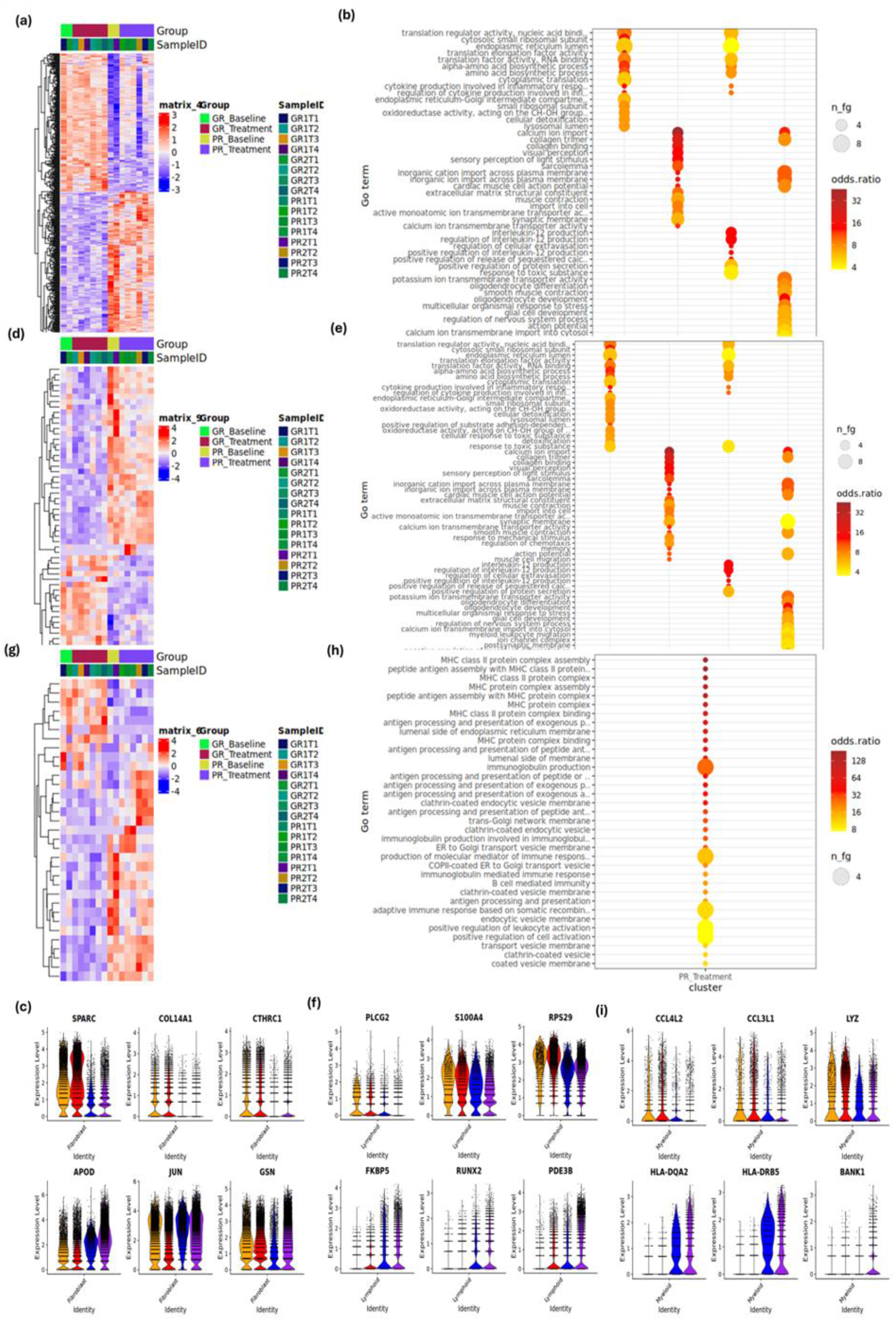
Pseudobulk differential expression analysis and functional analysis of skin cell subpopulations. (a) Heatmap of differential gene expression in fibroblast population between GR and PR. (b) GO Term enrichment analysis of biological process, molecular function and cellular components of DEGs in fibroblasts between GR and PR. (c) Violin plot illustrating the expression of some DEGs in fibroblast population between GR and PR. (d) Heatmap of differential gene expression in lymphoid cell population between GR and PR. (e) GO Term enrichment analysis of biological process, molecular function and cellular components of DEGs in lymphoid population between GR and PR. (f) Violin plot illustrating the expression of some DEGs in lymphoid population between GR and PR. (g) Heatmap of differential gene expression in myeloid cell population between GR and PR. (h) GO Term enrichment analysis of biological process, molecular function and cellular components of DEGs in myeloid population between GR and PR. (i) Violin plot illustrating the expression of some DEGs in myeloid population between GR and PR. GR1T1, Young scar 1, Baseline; GR1T2, Young scar 1, D21; GR1T3, Young scar 1, M6; GR1T4, Young scar 1, Endpoint (Y1); GR2T1, Young scar 2, Baseline; GR2T2, Young scar 2, D21; GR2T3, Young scar 2, M6; GR2T4, Young scar 1, Y1; PR1T1, Old scar 1, Baseline; PR1T2, Old scar 1, D21; PR1T3, Old scar 1, M6; PR1T4, Old scar 1, Endpoint (Y1); PR2T1, Old scar 2, Baseline; PR2T2, Old scar 2, D21; PR2T3, Old scar 2, M6; PR2T4, Old scar 1, Y1.

**Supplementary Figure 4.**
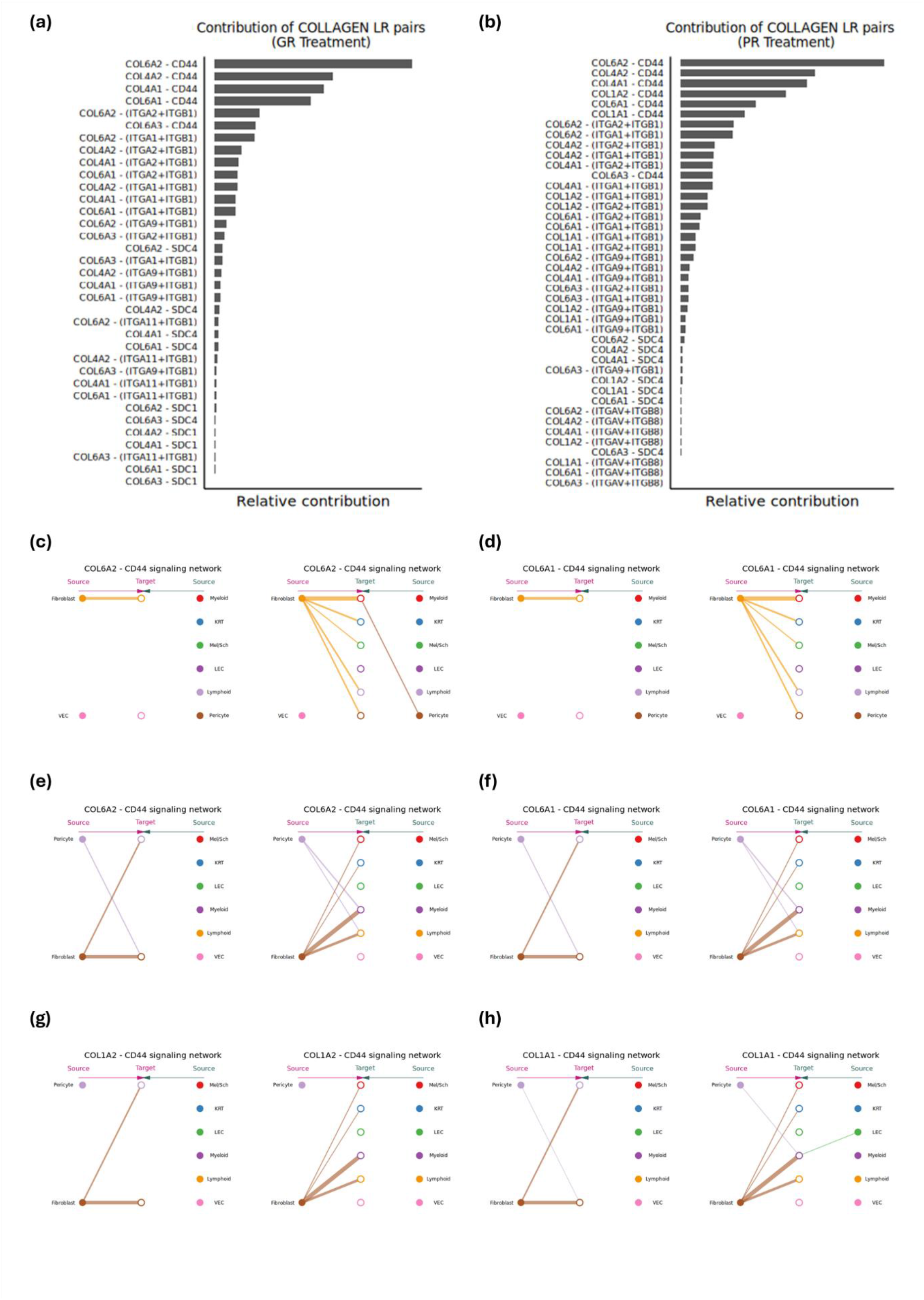
Intercellular collagen signalling networks in GR and PR. (a) Relative contribution of each ligand-receptor pairs in ECM collagen network in GRTx. (b) Relative contribution of each ligand-receptor pairs in ECM collagen network in PR. (c) Hierarchical plot showing inferred COL6A2-CD44 intercellular communication network in GRTx. (d) Hierarchical plot showing inferred COL6A1-CD44 intercellular communication network in GRTx. (e) Hierarchical plot showing inferred COL6A2-CD44 intercellular communication network in PRTx. (f) Hierarchical plot showing inferred COL6A1-CD44 intercellular communication network in PRTx. (g) Hierarchical plot showing inferred COL1A2-CD44 intercellular communication network in PRTx. (h) Hierarchical plot showing inferred COL1A1-CD44 intercellular communication network in PRTx.

**Supplementary Figure 5.**
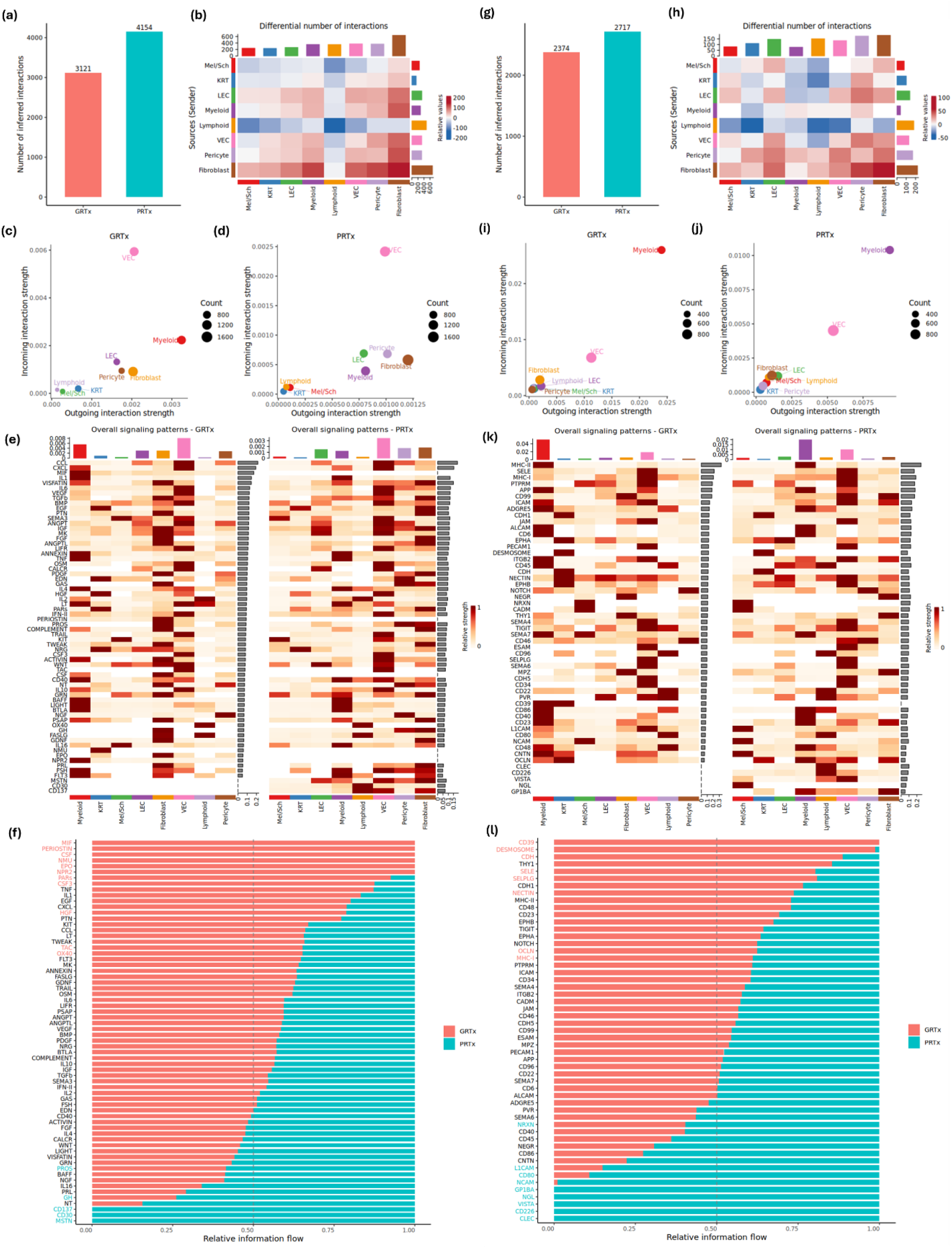
GR and PR scars have different patterns of secretion signalling and cell-cell intercellular communication networks. (a) Number of inferred interactions within secretion signalling communications networks in GRTx and PRTx. (b) Differential number of intercellular interactions in secretion signalling networks between GRTx and PRTx. Red, increased number of inter-cellular interactions in PRTx, compared to GRTx. Blue: decreased number of intercellular interactions in PRTx, compared to GRTx. Comparing the major sources and targets of SS network in (c) GRTx and (d) PRTx. (e) Overall secretion signalling patterns in GRTx (left panel) and PRTx (right panel). (f) Relative information flow of secretion signalling pathways in GRTx and PRTx. Red, GRTx; Green, PRTx. SS Pathways were ranked based on the overall information flow within the inferred networks between GRTx and PRTx. Significant pathways in GRTx were highlighted in Red on Y-axis and in PRTx were highlighted in Green on Y-axis. (g) Number of inferred interactions within cell-cell communications networks in GRTx and PRTx. (h) Differential number of intercellular interactions in CC networks between GRTx and PRTx. Red, increased number of inter-cellular interactions in PRTx, compared to GRTx. Blue: decreased number of intercellular interactions in PRTx, compared to GRTx. Comparing the major sources and targets of SS network in (i) GRTx and (j) PRTx. (k) Overall CC signalling patterns in GRTx (left panel) and PRTx (right panel). (l) Relative information flow of CC signalling pathways in GRTx and PRTx. Red, GRTx; Green, PRTx. SS Pathways were ranked based on the overall information flow within the inferred networks between GRTx and PRTx. Significant pathways in GRTx were highlighted in Red on Y-axis and in PRTx were highlighted in Green on Y-axis.

